# Accelerated Protein Biomarker Discovery from FFPE tissue samples using Single-shot, Short Gradient Microflow SWATH MS

**DOI:** 10.1101/675348

**Authors:** Rui Sun, Christie Hunter, Chen Chen, Weigang Ge, Nick Morrice, Shuang Liang, Chunhui Yuan, Qiushi Zhang, Xue Cai, Xiaoyan Yu, Lirong Chen, Shaozheng Dai, Zhongzhi Luan, Ruedi Aebersold, Yi Zhu, Tiannan Guo

## Abstract

We report and evaluated a microflow, single-shot, short gradient SWATH MS method intended to accelerate the discovery and verification of protein biomarkers in clinical specimens. The method uses 15-min gradient microflow-LC peptide separation, an optimized SWATH MS window configuration and OpenSWATH software for data analysis.

We applied the method to a cohort 204 of FFPE prostate tissue samples from 58 prostate cancer patients and 10 prostatic hyperplasia patients. Altogether we identified 27,976 proteotypic peptides and 4,043 SwissProt proteins from these 204 samples. Compared to a reference SWATH method with 2-hour gradient the accelerated method consumed only 27% instrument time, quantified 80% proteins and showed reduced batch effects. 3,800 proteins were quantified by both methods in two different instruments with relatively high consistency (r = 0.77). 75 proteins detected by the accelerated method with differential abundance between clinical groups were selected for further validation. A shortlist of 134 selected peptide precursors from the 75 proteins were analyzed using MRM-HR, exhibiting high quantitative consistency with the 15-min SWATH method (r = 0.89) in the same sample set. We further verified the capacity of these 75 proteins in separating benign and malignant tissues (AUC = 0.99) in an independent prostate cancer cohort (n=154).

Overall our data show that the single-shot short gradient microflow-LC SWATH MS method achieved about 4-fold acceleration of data acquisition with reduced batch effect and a moderate level of protein attrition compared to a standard SWATH acquisition method. Finally, the results showed comparable ability to separate clinical groups.

## INTRODUCTION

A large number of clinical and pre-clinical research questions require biomarkers for the classification of samples or phenotypes. Because they are thought to closely reflect the biochemical state of samples, protein biomarkers are particularly valuable. Protein biomarkers have been intensely sought to indicate disease type or stage, to report disease progression or response or resistance to treatment. For the most part protein biomarker projects use mass spectrometry as the base technique. In spite of enormous research efforts, the number of protein biomarkers discovered by proteomic methods that have progressed to clinical utility remains small (1-4).

Protein biomarker discovery and validation projects face significant technical and logistical challenges, including the following: i) biological protein abundance variability. Useful protein biomarkers will only be discovered if the variability within a population is smaller than the variability between protein groups. In the context of a twin cohort study of plasma proteins we have shown that the variability of proteins and the root cause for the variability varies greatly in a human population and that particularly variable proteins are unlikely to be selected as biomarkers (5). ii) confounding effects. Protein biomarker studies suffer from a range of confounding effects, including batch effects of sample collection, sample processing, data acquisition and data analysis. Batch effects are particularly severe among different cohorts that might be required to validate results from a discovery cohort, iii) sample availability. Frequently, sample cohorts of sufficient size and quality to generate sufficient statistical power are not available and iv) technical limitations. Even if suitable cohorts are available acquiring reproducible protein patterns by mass spectrometry from extended cohorts has been costly and challenging. For example, typically, protein biomarkers have multi-dimensional fractionation of the peptides generated from digested, tissue-extracted proteins followed by the analysis of the fractions by shotgun MS analysis. Even if isotopic or isobaric labeling methods increase the multiplexing capability of such analyses (6), the general approach remains expensive and technically challenging (7-9). Overall, these challenges convincingly support the need for the proteomic measurement of large sample cohorts at moderate cost, limited batch effects and high degree of reproducibility. At present state-of-the art, large scale clinical proteomic studies consist of 100 to 200 clinical samples (9-11) and there are indication, *e.g* the lack of stability of discovered marker panels that suggest that this number of samples is at the lower end of the required size range(12). Further, these studies were for the most part carried out by highly specialized groups or consortia using highly optimized analytical platforms. For many proteomic research groups that lack the means to implement the involved consortia methods, meaningful protein biomarker studies have therefore remained out of reach. Therefore, there is an urgent need for robust, highly reproducible, high throughput methods that support large-scale biomarker studies at moderate cost and with limited time consumption.

Sample throughput can be increased with short LC gradients for the separation of peptides. Bekker-Jensen et.al have combined multiple dimension pre-fractionation with relatively short LC gradient using shotgun proteomics to achieve deep proteome analysis (13); however, this approach lacks reproducibility and it is still time-consuming for a large cohort study. We and others have found that SWATH/DIA mass spectrometry (14) is a more suitable acquisition method to classify samples in large sample cohorts(5, 15-17). SWATH/DIA is an acquisition method for biomarker studies because it identifies and quantifies peptide precursors via peak groups consisting of fragment ion chromatograms from highly convoluted mass spectra (15) and thus obviates the need to isolate peptide precursors during acquisition. This improves data completeness and enables efficient single-shot proteomic analysis. The key to this MS technique is the ability to collect high-resolution MS/MS spectra at very high acquisition rates, such that a wide mass range can be covered with a series of smaller Q1 isolation windows in an LC compatible cycle time. Thus the fast scanning rate of TripleTOF system has been the key in enabling the shortening of LC gradients for analyzing complex tissue proteomes, from 120 min (15) to 45 min (17) without strongly compromising proteome depth, and has been increasingly applied to analyze various types of clinical samples including plasma (5) and tumor tissues (15, 17, 18). A faster nano-LC and Orbitrap-based MS method has been reported recently to allow analysis of plasma and cell line samples using a 21-min gradient (19). However, this method requires specialized LC system.

To further improve the robustness and throughput of the proteomic analysis of sizable sample cohorts, the use of microflow chromatography is a promising option. An increasing number of studies have demonstrated the applicability of microflow coupled with SWATH MS (20-24). E.g. the Ralser group applied microflow-LC and SWATH to study yeast proteome at a throughput of 60 samples per day (24).

Here, we established and optimized a 15-min gradient microflow LC SWATH method, and rigorously examined its performance by analyzing 204 FFPE prostate tissue samples. From the detected 4,043 proteins we prioritized 75 that were further verified with respect to their ability to separate cancer from hyperplasia in an independent FFPE prostate tissue sample cohort study by complementary methods. The results indicate the that short gradient microflow-LC SWATH is a suitable and robust method for clinical protein biomarker studies.

## EXPERIMENTAL PROCEDURES

### Standard protein digests

Digests of proteins isolated from HEK 293 cell were prepared as previously described (25) and provided by Dr Yansheng Liu from ETH Zurich (now in Yale University). Protein digests from K562 cells were obtained from the SWATH Performance Kit (SCIEX). 10% (v/v) iRT peptides (Biognosys, Switzerland) were spiked into peptide samples prior to MS analysis for retention time calibration.

### PCa patient cohorts and formalin-fixed paraffin-embedded (FFPE) samples

Two prostate cancer (PCa) sample cohorts termed PCZA and PCZB were used in this study. The PCZA was acquired by the Second Affiliated Hospital College of Medicine, Zhejiang University and consisted of 58 PCa patients and 10 benign prostatic hyperplasia (BPH) patients. The PCZB cohort was acquired by the Second Affiliated Hospital College of Medicine, Zhejiang University and consisted of 24 PCa patients and 30 BPH patients whose benign and hyperplastic regions have been distinguished. All patients were recruited in 2017 and 2018. All cohorts were approved by the ethics committee of the respective hospitals for the procedures of this study.

The two different cohort samples were handled by different pathology laboratories, fixed and embedded by the respective staff. The samples were similarly processed and analyzed at different time points. For the PCZA cohort, three biological replicates (size 1 × 1 × 5 mm^3^) were collected and analyzed by SWATH MS and MRM-HR. For the PCZB cohort, two biological replicates (1.5 × 1.5 × 5 mm^3^) were analyzed by MRM-HR.

### Pressure cycling technology (PCT)-assisted peptide extraction from FFPE tissues

About 0.5 mg of FFPE tissue was punched from the samples, weighed and processed for each biological replicate via the FFPE-PCT workflow as described previously (26). Briefly, the tissue punches were first dewaxed by incubating with 1 mL of heptane under gentle vortexing at 600–800 rpm, followed by serial rehydration using 1 mL of 100%, 90%, and 75% ethanol (General reagent, G73537B, Shanghai, China), respectively. The samples were further incubated with 200 μL of 0.1% formic acid (FA) (Thermo Fisher Scientific, T-27563) at 30 °C for 30 min for acidic hydrolysis. The tissue punches were then transferred into PCT-MicroTubes (Pressure Biosciences Inc., Boston, MA, USA, MT-96) and briefly washed with 100 μL of freshly prepared 0.1 M Tris-HCl (pH 10.0) to remove residual FA. Thereafter, the tissues were incubated with 15 μL of freshly prepared 0.1 M Tris-HCl (pH 10.0) at 95 °C for 30 m in with gentle vortexing at 600 rpm. Samples were immediately cooled to 4 °C after basic hydrolysis.

Following the pretreatment described above, 25 μL of lysis buffer including 6 M urea (Sigma, U1230), 2 M thiourea (Amresco, M226) in 100 mM ammonium bicarbonate (General regent, G12990A, Shanghai, China), pH 8.5 was added to the PCT-MicroTubes containing tissues. The tissue samples were further subjected to PCT-assisted tissue lysis and protein digestion procedures using the Barocycler NEP2320-45K (Pressure Biosciences Inc., Boston, MA, USA) as described previously (27). The PCT scheme for tissue lysis was set with each cycle containing a period of 30 s of high pressure at 45 kpsi and 10 s at ambient pressure, oscillating for 90 cycles at 30°C. Protein reduction and alkylation was performed at ambient pressure by incubating protein extracts with 10 mM Tris(2-carboxyethyl) phosphine (TCEP) (Sigma, C4706) and 20 mM iodoacetamide (IAA) (Sigma, I6125) in darkness at 25 °C for 30 min, with gentle vortexing at 600 rpm in a thermomixer. Then the proteins were digested with MS grade Lys-C (Hualishi, Beijing, China, enzyme-to-substrate ratio, 1:40) using a PCT scheme with 50 s of high pressure at 20 kpsi and 10 s of ambient pressure for each cycle, oscillating for 45 cycles at 30 °C. Thereafter, the proteins were further digested with MS grade trypsin (Hualishi, Beijing, China, enzyme-to-substrate ratio, 1:50) using a PCT scheme with 50 s of high pressure at 20 kpsi and 10 s of ambient pressure in one cycle, oscillating for 90 cycles at 30 °C. Peptide digests were then acidified with 1% trifluoroacetic (TFA) (Thermo Fisher Scientific, T/3258/PB05) to pH 2–3 and subjected to C18 desalting. iRT peptides were spiked into peptide samples at a final concentration of 10% prior to MS analysis for RT calibration.

### Optimization of microflow LC gradients coupled with SWATH MS

During the optimization studies, 1 μg peptides were separated with different microflow gradients and different SWATH MS parameters. Linear gradients of 3–35% acetonitrile (0.1% formic acid) with durations of 5, 10, 20, 30, and 45 min were evaluated. The number of Q1 variable windows (40, 60, 100) and MS/MS accumulation times (15, 25 ms) constituted the key parameters that were adjusted for the shorter gradients. The need for collision energy spread with the optimized collision energy ramps was tested. Four replicates were performed for each test, after which the data were processed with the PeakView® software with the SWATH 2.0 MicroApp to evaluate the number of proteins and peptides quantified with FDR < 1 % and CV < 20%. The optimized methods were then tested on multiple instruments with different cell lysates to confirm the robustness of the method.

### SWATH MS acquisition

Peptides were separated at a flow rate of 5 μL/min by a 15-min SWATH of 5–35% linear LC gradient elution (buffer A: 2% ACN (Sigma, 34851), 0.1% formic acid; buffer B: 80% ACN, 0.1% formic acid) on a column, 3 μm, ChromXP C18CL, 120 Å, 150 x 0.3 mm using an Eksigent NanoLC™ 400 System coupled with a TripleTOF^®^ 6600 system (SCIEX). The DuoSpray Source was replumbed using the 25 μm ID hybrid electrodes to minimize post-column dead volume. The applied SWATH method was composed of a 150 ms TOF MS scan with m/z ranging from 350 to 1250 Da, followed by MS/MS scans performed on all precursors (from 100 to 1500 Da) in a cyclic manner. A 100 variable Q1 isolation window scheme was used in this study (Supplementary Table 1B). The accumulation time was set at 25 ms per isolation window, resulting in a total cycle time of 2.7 s.

We also included beta-galactosidase digest (β-gal) (SCIEX, 4465867) for mass and retention time calibration which was analyzed every four injections. The target ion (m/z = 729.4) which is from a peptide precursor in the β-gal digest mixture was monitored under high sensitivity mode. The RT, intensity, and m/z of targeted precursor and fragment ions were respectively used for LC QC, the sensitivity test, and mass calibration separately.

### MRM-HR MS acquisition

A time scheduled MRM-HR targeted quantification strategy was used to further validate proteins observed to be differentially expressed proteins by SWATH MS as described above. The same microflow LC approach was used for 15-min SWATH MS analysis. The TripleTOF 6600 mass spectrometer was operated in IDA mode for time-scheduling the MS/MS acquisition for 134 peptides for the MRM-HR workflow. The method consisted of one 75 ms TOF-MS scan for precursor ions with m/z ranging from 350 to 1250 Da, followed by MS/MS scans for fragment ions with m/z ranging from 100 to 1500 Da, allowing for a maximum of 45 candidate ions being monitored per cycle (25 ms accumulation time, 50 ppm mass tolerance, rolling collision energy, +2 to +5 charge states with intensity criteria above 2 000 000 cps to guarantee that no untargeted peptides should be acquired). The fragment ion information including m/z and RT of a targeted precursor ion was confirmed by previous SWATH results and was then added to the inclusion list for the targeted analysis. The intensity threshold of targeted precursors in the inclusion list was set to 0 cps and the scheduling window was 60 s. The targeted peptide sequences were the same as those found in the previous SWATH MS analysis.

Targeted MRM-HR data were analyzed by 19.0.9.149 Skyline (28), which automatically detected the extracted-ion chromatogram (XIC) from an LC run by matching the MS spectra of the targeted ion against its spectral library generated from the IDA mode within a specific mass tolerance window around its m/z. All peaks selected were checked manually after automated peak detection using Skyline. Both MS1 and MS2 filtering were set as “TOF mass analyzer” with a resolution power of 30 000 and 15 000, respectively, while the “Targeted” acquisition method was defined in the MS/MS filtering.

### SWATH data analysis

The optimization data for optimal LC gradients were processed using the SWATH 2.0 MicroApp in PeakView^®^ software (SCIEX) using the pan-human library (29). RT calibration was performed by first using iRT peptides with RT window at a 75 ppm XIC extraction width. Replicate analysis was performed using the SWATH Replicate Analysis Template (SCIEX) to determine the number of peptides and proteins quantified with FDR < 1% peptide and CV < 10 or 20%.

The data from prostate samples were processed using the OpenSWATH pipeline. Briefly, SWATH raw data files were converted in profile mode to mzXML using msconvert and analyzed using OpenSWATH (2.0.0) (30) as described previously (15). The retention time extraction window was 600 s, while m/z extraction was performed with 0.03 Da tolerance. RT was then calibrated using both iRT peptides. Peptide precursors were identified by OpenSWATH and PyProphet (2.0.1) with d_score < 0.01 and FDR < 1%. For each protein, the median MS2 intensity value of peptide precursor fragments which were detected to belong to the protein was used to represent the protein abundance.

## RESULTS AND DISCUSSIONS

### Establishment and optimization of the 15-min microflow SWATH MS method

A HEK 293 cell lysate digest was used to establish and optimize the short microflow LC gradient and SWATH acquisition schemes on TripleTOF 6600 systems. Specifically, we tested the effects of LC gradient lengths of 5, 10, 20, and 45 min, and mass spectrometer parameters including variable Q1 windows and accumulation time for MS2 (Supplementary Table 1). For each injection 1 μg mass of total peptide was loaded onto a microflow column of 150 x 0.3 mm dimensions and analyzed under a range of conditions. To increase robustness of results, four technical replicates of each condition were used. The acquired data were searched consistently searched against PHL with the PeakView® software and the SWATH 2.0 MicroApp and the number of peptides and inferred proteins, as well as their intensities were recorded. The data was processed as described in the methods section and evaluated according to the number of proteins and peptides identified with FDR < 1% and quantified with CV < 10% or CV < 20%, respectively. The whole dataset was acquired on two different instruments. Supplementary Figure 1a shows that using shorter gradient methods generated similar results between the two different 6600 instruments. Our data also showed that the 20 min microflow method detected 90% of the proteins quantified by 45 min method, while the 10 min LC method identified 70% of the proteins. With decreasing gradient length, the number of identified proteins further decreased to 77% for a 10 min method to 53% for a 5 min method in their best condition (Supplementary Figure 1a).

Next, we optimized the specific mass spec parameters including variable windows and accumulation times to balance the width of the windows and scan times (Supplementary Figure 1b). Typically, more variable windows led to more peptides and proteins quantified robustly, but only up to a point where the MS/MS acquisition rates become too fast or the cycle times too long, as evidenced in the 5min gradient optimization results. Thus, a higher number of variable windows led to a higher number of peptide and protein identifications. The optimal accumulation time was highly dependent on the LC time. Higher numbers of acquisition windows necessitated shorter MS/MS accumulation times per precursor ion window to maintain a cycle time that was compatible with the peak width generated by the respective gradients. Considering the tradeoff between sample throughput and numbers of proteins quantified, the gradient time from 10 min to 20 min is a better choice according to the efficiency of peptides and proteins identification in unit of time (Supplementary Figure 1b). Therefore, we chose the 15 min gradient as the optimal LC condition (Supplementary Figure 2).

### Application of short gradient microflow-SWATH to a PCa patient cohort

We evaluated the performance of the optimized short gradient microflow LC SWATH method on a set of prostate cancer (PCa) tissue samples named PCZA. The set consisted of 204 FFPE biospecimens collected from 58 PCa patients and 10 benign hyperplasia (BPH) patients (Supplementary Table 2) for which clinical data were also available. The 204 samples were randomly divided into seven batches and digested into peptides in barocyclers. Every batch included a mouse liver sample as quality control (QC) for the PCT-assisted sample preparation and a prostate tissue pool samples as the QC sample for SWATH MS (Supplementary Figure 3).

We then subjected the resulting peptide samples to the15-min-SWATH method optimized above (Figure 1a). The total sample set consisted of 58 PCa samples and 7 QC and reference samples. The 204 samples were measured in 125.7 hrs (∼5 days) and quantified 27,975 peptide precursors from 4,038 SwissProt proteins (without protein grouping) with 74.79% missing value rate. On average, 5,615 peptide precursors from 1,018 proteins were quantified for each sample. More peptides and proteins were quantified from tumor samples (5,861 peptide precursors from 1,078 proteins on average) than benign samples (3,988 peptide precursors from 618 proteins on average). Totally 913 proteins were quantified in at least 50% samples (Supplementary Table 3).

**Figure 1.**
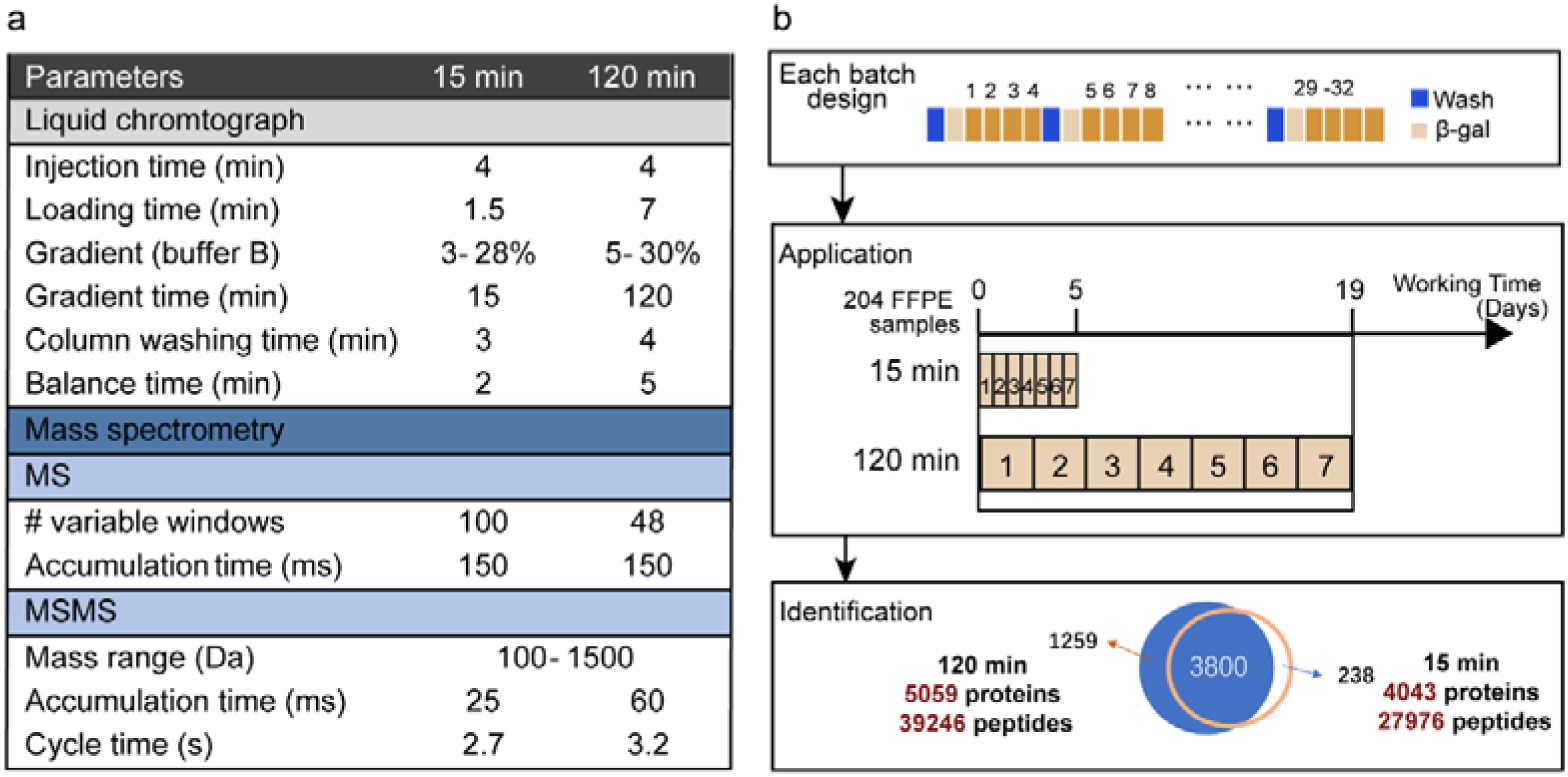
The short-gradient SWATH method and application in the PCZA PCa cohort. (a) Comparison of parameters between the 15-min SWATH and 120-min SWATH methods. (b) The workflow of the 15-min-SWATH and conventional SWATH for the PCZA cohort. We designed seven randomly shuffled batches with a column washing run and a calibration (β-gal) run inserted every four samples.

To allow a comparison of the accelerated short gradient method with a standard SWATH MS method with respect to the number of proteins recorded and the respective clinically relevant information content we re-acquired the whole sample set with a 120-min LC gradient and 48 variable Q1 windows in a TripleTOF 5600+(26). These measurements consumed 467 hr (∼20 days) and identified 38,338 peptide precursors from 5,059 SwissProt proteins with 61.86% missing value rate. On average, 10,751 peptide precursors from 1,921 proteins were quantified for each sample. More peptides and proteins were quantified from tumor samples (11,439 peptide precursors from 2,054 proteins on average) than benign samples (6,693 peptide precursors from 1192 proteins on average). Totally 1,914 proteins were quantified in at least 50% samples (Supplementary Table 3). Compare to this 120-min method, the 15-min method characterized about half of peptide precursors and proteins.

Overall, the data shows that the 15-min-SWATH coverage reached 50-80% of that achieved by a standard method. In all samples, 3,800 proteins were quantified by both methods. This result was generated at a 6-fold reduced acquisition time (time 125.7 hrs vs, 467 hrs) (Figure 1b) suggesting that clinical cohorts of significant size can be measured by the accelerated method quickly, efficiently.

### Reproducibility and batch effect analysis

We evaluated the reproducibility of the datasets produced by the 15-min gradient and the 120-min gradient SWATH with respect to reproducibility and batch effect. We first assessed the technical reproducibility by correlation between technical replicates for LC-MS. The technical reproducibility of the data obtained by the 15-min SWATH method (r = 0.99) is slightly higher than that from the 120-min SWATH method (r = 0.86) (Figure 2a). Thus, the measured biological reproducibility is also slightly higher in the 15-min SWATH method (Figure 2a). If we focused the analysis on the 3,800 proteins quantified by both methods, we observed a high degree of similarity (r = 0.7681) between the methods (Figure 2b).

**Figure 2.**
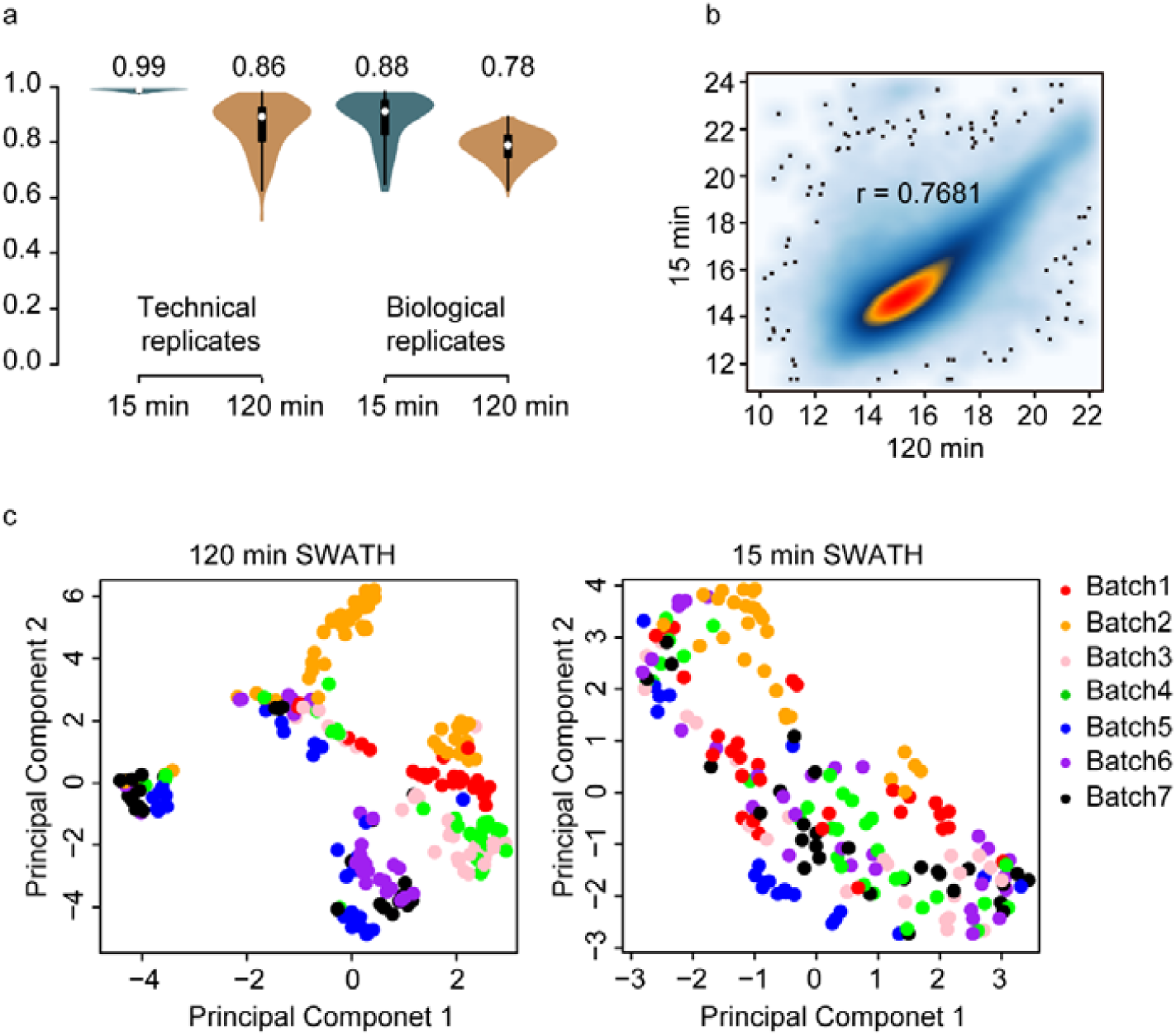
The reproducibility of the short gradient SWATH method in the PCZA PCa cohort. (a) Violin plots show the technical replicates and biological replicates in the two methods. (b) Pearson correlation of log2-scaled protein intensity values obtained from 3,800 proteins that were quantified by both methods. (c) PCA analysis of all samples quantified by the 120-min method (left) and the 15-min method (right).

We next analyzed batch effects apparent in either dataset. Batch effects are an unavoidable reality resulting from technical variation in multi-day MS analyses and are a non-trivial complication for big cohort proteomics analysis. Several algorithms have been developed to bioinformatically minimize the missing value rate, however, these imputation approaches remain controversial (31). We evaluated the batch effect of the data acquired by the 15-min SWATH, which is lower than that from the 120-min method (Figure 2c). Together, the 15-min SWATH method improved quantitative reproducibility and reduced batch effect.

### Verification of differential expression proteins using MRM-HR

On the path to clinical or preclinical use protein biomarkers detected by MS based cohort studies face a number of verification and validation requirements. These include technical verification of the abundance changes detected in the cohort study and validation in independent sample cohorts.

To further validation the abundance changes detected in the SWATH data we selected a panel of 75 proteins showing different abundance (absolute fold change larger than two and adjusted p-value less than 0.05) between control and tumor tissue and measured their respective intensities using the targeted MS method MRM-HR. The selected proteins were associated with most strongly cancer dis regulated pathways and included 21 known diagnosis biomarkers such as ACPP and FASN, and 10 drug targets (Supplementary Table 4A). The proteins were further annotated in IPA (Supplementary Table 4B) indicating that the proteins suggested elevated cell migration, development and growth, and suppressed cell death and survival (Supplementary Figure 5).

For these measurements the MRM-HR method was optimized using a pooled prostate sample to determine the best performing peptides from the selected proteins, and best target fragment ions for quantitation. The information about proteins and peptides including the RTs was imported into Skyline to build a spectral library. A total of 134 peptides for 75 proteins were selected for targeted detection (Supplementary Table 2E). Time scheduling was used to ensure at least eight data points were obtained across the LC peaks as well as an optimized accumulation time of 25 ms for each peptide for high-quality quantitative data.

To confirm the quantitative accuracy of the 15-min SWATH data, we re-analyzed 99 samples in the PCZA cohort using the MRM-HR method. The 99 samples were randomly allocated to five batches, each containing 20 samples and an extra MS QC sample which was a pool of prostate tissue digests in PCZA. We firstly examined the reproducibility of XICs for all peptides in MRM-HR assays. For the five pooled samples measured across five batches, we found that 76.6% of precursors measured from the peptides were quantified with a CV below 20%. The median CV was 13.4% (Supplementary Figure 6). Next the protein fold-changes between tumor and normal samples were calculated to investigate the correlation of 15-min SWATH with MRM-HR (Figure 3a).

**Figure 3.**
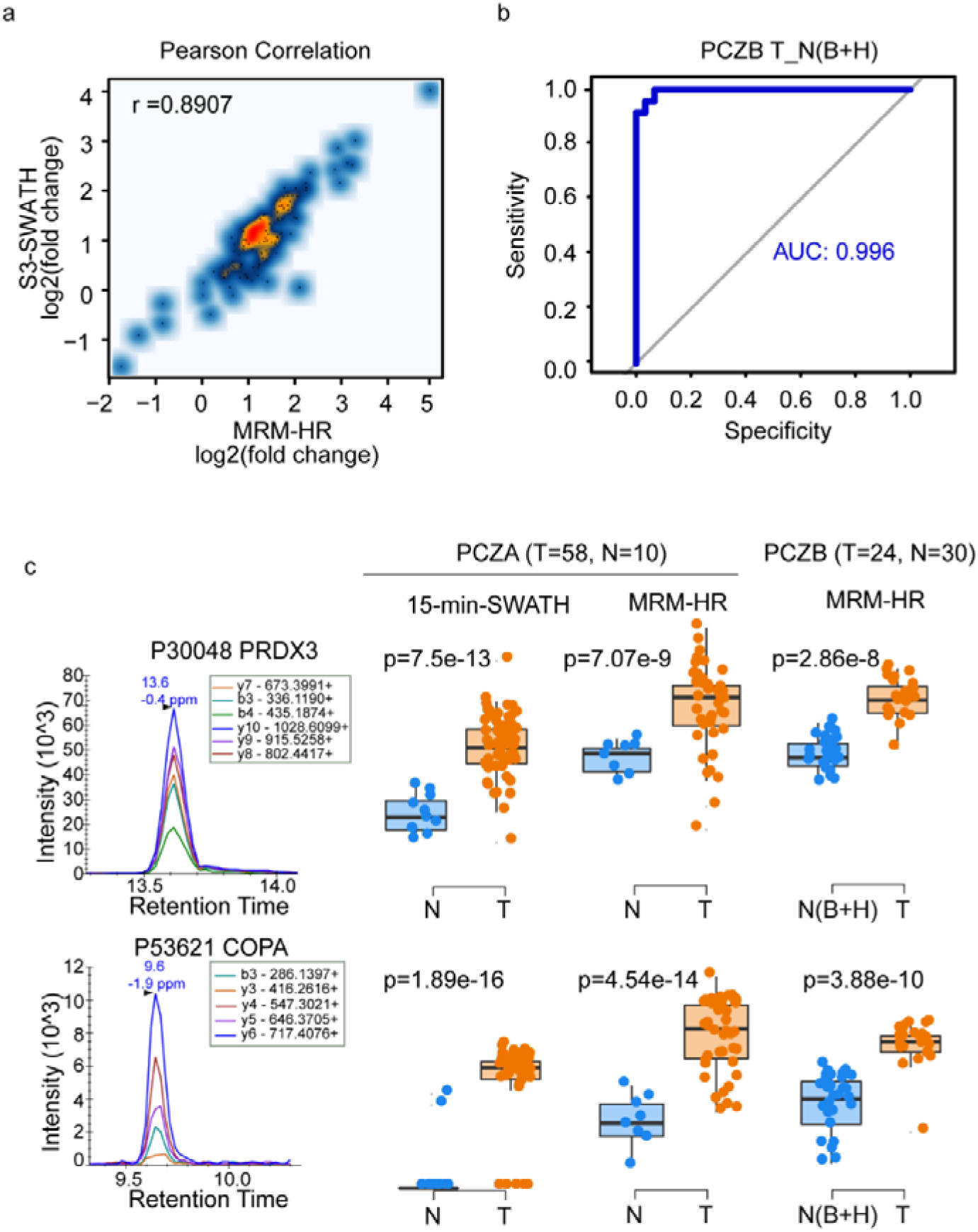
Verification of proteomic data using MRM-HR. (a) Pearson correlation coefficient between the 15-min SWATH and MRM-HR datasets based on the log_2_(T/N) of protein expression in PCZA. (b) The ROC curves of protein quantification from MRM-HR to predict the tumor and normal tissues with the random forest algorithm in PCZB (T: PCa, N: BPH, H: hyperplasia in BPH patients, B: benign in BPH patients). (c) MRM-HR validation of potential diagnostic proteins using the PCZA and PCZB. PRDX3 (peptide: +2 DYGVLLEGSGLALR), COPA (peptide: +2 DVAVMQLR). The left panel shows the fragment ion extracted-ion chromatograms (XICs) for the peptide from each protein. The right panel of boxplots shows the peptides quantified in the different data sets.

We further quantified the expression levels of the 75-protein-panel in an independent prostate cancer cohort, PCZB, containing 30 BPH and 24 PCa in duplicated biological replicates using the same 15-min SWATH MRM-HR workflow (Supplementary Table 5). For the six pooled samples measured across six batches, 75.6% of peptide precursors were quantified with a CV below 20%. The median CV is 14.9% (Supplementary Figure 6).

To assess the power of the protein panel of differentially abundant proteins to separate benign and malignant tissues, we assembled a random-forest model for the PCZA MRM-HR dataset, and found an accuracy of 0.992 in this set (Supplementary Figure 7). Next, we tested the ability of this panel to separate tumor from benign prostatic tissue samples in an independent patient group, *i.e.* PCZB, including 24 PCa patients and 30 BPH patients. The receiver operating curves (ROC) of the 75-protein-panel clearly distinguished PCa from BPH patient groups (Figure 3b).

We then investigated in detail two proteins—PRDX3 (P30048) and COPA (P53621) which were prioritized because of their role in TP53 oncogene regulation and as a potential drug (decitabine) target (Supplementary Figure 8). The data show that these proteins significantly up-regulated in tumor tissue from all three workflows, *i.e.*15-min-SWATH, and MRM-HR in the PCZA cohort samples and MRM-HR in the PCZB cohort (Figure 3b). The ROC curve of these two proteins from three different datasets distinguishing benign from malignant tissue samples are shown in Supplementary Figure 9, with all of AUC over 0.78. Taken together, we validated these dysregulated proteins quantification by SWATH showed higher reliability and performed better prediction ability in different sample cohorts.

### Conclusion

In this study, we present a 15-min microflow-LC SWATH that supports the consistent proteomic analysis of clinical (FFPE) samples at a throughput of ∼50 samples per day (excluding calibration and washing). The method is therefore well suited for the analysis of large sample cohorts, even in a single investigator proteomic laboratory. The results show that the presented method increases the throughput by ca six-fold compared to a conventional SWATH MS method, at reduced batch effects and at an attrition of ca 20% of detected proteins and increased missing value rate (∼20% worse) in the prostate cancer cohort. For individual samples, the number of detected proteins decreased by ∼50%. The quantitative accuracy of the short gradient method was comparable to that achieved by targeted quantification using MRM-HR for shortlisted proteins. This work showed the potential of this short gradient SWATH proteomics pipeline for accelerated discovery and verification of protein biomarkers for precision medicine.

## Author Contributions

T.G., C.H., R.S. designed the project. C.H., N.M. and C.C. optimized the 15-min-SWATH. X.Y., L.C. procured the three prostate cohorts. R.S. performed the PCT SWATH analysis with help from X.C. C.C. and R.S. performed the MRM-HR analysis. W.G., R.S., S.D., analyzed the data. R.S., Y.Z., C.H., R.A. and T.G. wrote the manuscript. Z.L assisted data analysis. S.L., C.Y. gave valuable advice. T.G., Y.Z. supported and supervised the project.

## Research Funding

Zhejiang Provincial Natural Science Foundation of China (Grant No. LR19C050001 to T.G.). Hangzhou Agriculture and Society Advancement Program (Grant No. 20190101A04 to T.G.). National Natural Science Foundation of China (General Program) (Grant No. 81972492 to T.G.). National Science Fund for Young Scholars (Grant No. 21904107).

## Acknowledgments

We thank Dr Xuan Ding for review of the manuscript.

## Competing financial interests

The research group of T.G. is partly supported by SCIEX, which provides access to prototype instrumentation, and Pressure Biosciences Inc, which provides access to advanced sample preparation instrumentation.

## Data and materials availability

The 15-min SWATH data are deposited in PRIDE. Project accession: IPX0001645000. The 15-min SWATH data are deposited in iProX (IPX0001645001). The MRM-HR data are deposited in iProX (IPX0001645002). All the data will be publicly released upon publication.

## ABBREVIATIONS

AUC: area under the curve
BPH: benign prostatic hyperplasia
CV: coefficient of variation
DDA: data dependent acquisition
DIA: data independent acquisition
FA: formic acid
FDR: false discovery rate
FFPE: formalin fixed, paraffin embedded
IAA: iodoacetamide
IPA: ingenuity pathway analysis
LC: liquid chromatograph
MRM-HR: multiple reaction monitoring high-resolution
PCa: prostate cancer
PCT: pressure cycling technology
PRM: parallel reaction monitoring
QC: quality control
ROC: receiver operating characteristic
RF: random forest
RT: retention time
SWATH MS: sequential windowed acquisition of all theoretical fragment ion - mass spectra
TCEP: tris(2-carboxyethyl) phosphine
TFA: trifluoroacetic
TMA: tissue microarray analysis
TOF: time of flight
XIC: extracted ion chromatogram

## References

1. Olsen, M.; Ghannad, M.; Lok, C.; Bossuyt, P. M., Shortcomings in the evaluation of biomarkers in ovarian cancer: a systematic review. Clin Chem Lab Med 2019.

2. Rifai, N.; Gillette, M. A.; Carr, S. A., Protein biomarker discovery and validation: the long and uncertain path to clinical utility. Nat Biotechnol 2006, 24, (8), 971–83.

3. Anderson, N. L.; Ptolemy, A. S.; Rifai, N., The riddle of protein diagnostics: future bleak or bright? Clin Chem 2013, 59, (1), 194–7.

4. Frantzi, M.; Latosinska, A.; Kontostathi, G.; Mischak, H., Clinical Proteomics: Closing the Gap from Discovery to Implementation. Proteomics 2018, 18, (14), e1700463.

5. Liu, Y.; Buil, A.; Collins, B. C.; Gillet, L. C.; Blum, L. C.; Cheng, L. Y.; Vitek, O.; Mouritsen, J.; Lachance, G.; Spector, T. D.; Dermitzakis, E. T.; Aebersold, R., Quantitative variability of 342 plasma proteins in a human twin population. Mol Syst Biol 2015, 11, (1), 786.

6. Aebersold, R.; Mann, M., Mass-spectrometric exploration of proteome structure and function. Nature 2016, 537, (7620), 347–55.

7. Sabrkhany, S.; Kuijpers, M. J. E.; Knol, J. C.; Olde Damink, S. W. M.; Dingemans, A. C.; Verheul, H. M.; Piersma, S. R.; Pham, T. V.; Griffioen, A. W.; Oude Egbrink, M. G. A.; Jimenez, C. R., Exploration of the platelet proteome in patients with early-stage cancer. J Proteomics 2018, 177, 65–74.

8. Mun, D. G.; Bhin, J.; Kim, S.; Kim, H.; Jung, J. H.; Jung, Y.; Jang, Y. E.; Park, J. M.; Kim, H.; Jung, Y.; Lee, H.; Bae, J.; Back, S.; Kim, S. J.; Kim, J.; Park, H.; Li, H.; Hwang, K. B.; Park, Y. S.; Yook, J. H.; Kim, B. S.; Kwon, S. Y.; Ryu, S. W.; Park, D. Y.; Jeon, T. Y.; Kim, D. H.; Lee, J. H.; Han, S. U.; Song, K. S.; Park, D.; Park, J. W.; Rodriguez, H.; Kim, J.; Lee, H.; Kim, K. P.; Yang, E. G.; Kim, H. K.; Paek, E.; Lee, S.; Lee, S. W.; Hwang, D., Proteogenomic Characterization of Human Early-Onset Gastric Cancer. Cancer Cell 2019, 35, (1), 111–124 e10.

9. Zhang, H.; Liu, T.; Zhang, Z.; Payne, S. H.; Zhang, B.; McDermott, J. E.; Zhou, J. Y.; Petyuk, V. A.; Chen, L.; Ray, D.; Sun, S.; Yang, F.; Chen, L.; Wang, J.; Shah, P.; Cha, S. W.; Aiyetan, P.; Woo, S.; Tian, Y.; Gritsenko, M. A.; Clauss, T. R.; Choi, C.; Monroe, M. E.; Thomas, S.; Nie, S.; Wu, C.; Moore, R. J.; Yu, K. H.; Tabb, D. L.; Fenyo, D.; Bafna, V.; Wang, Y.; Rodriguez, H.; Boja, E. S.; Hiltke, T.; Rivers, R. C.; Sokoll, L.; Zhu, H.; Shih, I. M.; Cope, L.; Pandey, A.; Zhang, B.; Snyder, M. P.; Levine, D. A.; Smith, R. D.; Chan, D. W.; Rodland, K. D.; Investigators, C., Integrated Proteogenomic Characterization of Human High-Grade Serous Ovarian Cancer. Cell 2016, 166, (3), 755–765.

10. Vasaikar, S.; Huang, C.; Wang, X.; Petyuk, V. A.; Savage, S. R.; Wen, B.; Dou, Y.; Zhang, Y.; Shi, Z.; Arshad, O. A.; Gritsenko, M. A.; Zimmerman, L. J.; McDermott, J. E.; Clauss, T. R.; Moore, R. J.; Zhao, R.; Monroe, M. E.; Wang, Y. T.; Chambers, M. C.; Slebos, R. J. C.; Lau, K. S.; Mo, Q.; Ding, L.; Ellis, M.; Thiagarajan, M.; Kinsinger, C. R.; Rodriguez, H.; Smith, R. D.; Rodland, K. D.; Liebler, D. C.; Liu, T.; Zhang, B.; Clinical Proteomic Tumor Analysis, C., Proteogenomic Analysis of Human Colon Cancer Reveals New Therapeutic Opportunities. Cell 2019, 177, (4), 1035–1049 e19.

11. Mertins, P.; Mani, D. R.; Ruggles, K. V.; Gillette, M. A.; Clauser, K. R.; Wang, P.; Wang, X.; Qiao, J. W.; Cao, S.; Petralia, F.; Kawaler, E.; Mundt, F.; Krug, K.; Tu, Z.; Lei, J. T.; Gatza, M. L.; Wilkerson, M.; Perou, C. M.; Yellapantula, V.; Huang, K. L.; Lin, C.; McLellan, M. D.; Yan, P.; Davies, S. R.; Townsend, R. R.; Skates, S. J.; Wang, J.; Zhang, B.; Kinsinger, C. R.; Mesri, M.; Rodriguez, H.; Ding, L.; Paulovich, A. G.; Fenyo, D.; Ellis, M. J.; Carr, S. A.; Nci, C., Proteogenomics connects somatic mutations to signalling in breast cancer. Nature 2016, 534, (7605), 55–62.

12. Thomas. S.; Friedrich, B.,; Schnaubelt M.; Chan. D.; Zhang H,; Aebersold. R., Orthogonal proteomic platforms and their implications for the stable classification of high-grade serous ovarian cancer subtypes. BioRxiv 2019.

13. Bekker-Jensen, D. B.; Kelstrup, C. D.; Batth, T. S.; Larsen, S. C.; Haldrup, C.; Bramsen, J. B.; Sorensen, K. D.; Hoyer, S.; Orntoft, T. F.; Andersen, C. L.; Nielsen, M. L.; Olsen, J. V., An Optimized Shotgun Strategy for the Rapid Generation of Comprehensive Human Proteomes. Cell Syst 2017, 4, (6), 587–599 e4.

14. Gillet LC N. P., Tate S, Röst H, Selevsek N, Reiter L, Bonner R, Aebersold R., Targeted data extraction of the MS/MS spectra generated by data-independent acquisition: a new concept for consistent and accurate proteome analysis. Mol Cell Proteomics. 2012.

15. Guo, T.; Kouvonen, P.; Koh, C. C.; Gillet, L. C.; Wolski, W. E.; Rost, H. L.; Rosenberger, G.; Collins, B. C.; Blum, L. C.; Gillessen, S.; Joerger, M.; Jochum, W.; Aebersold, R., Rapid mass spectrometric conversion of tissue biopsy samples into permanent quantitative digital proteome maps. Nat Med 2015, 21, (4), 407–13.

16. Bouchal, P.; Schubert, O. T.; Faktor, J.; Capkova, L.; Imrichova, H.; Zoufalova, K.; Paralova, V.; Hrstka, R.; Liu, Y.; Ebhardt, H. A.; Budinska, E.; Nenutil, R.; Aebersold, R., Breast Cancer Classification Based on Proteotypes Obtained by SWATH Mass Spectrometry. Cell Rep 2019, 28, (3), 832–843 e7.

17. Zhu, Y.; Zhu, J.; Lu, C.; Zhang, Q.; Xie, W.; Sun, P.; Dong, X.; Yue, L.; Sun, Y.; Yi, X.; Zhu, T.; Ruan, G.; Aebersold, R.; Huang, S.; Guo, T., Identification of Protein Abundance Changes in Hepatocellular Carcinoma Tissues Using PCT-SWATH. Proteomics Clin Appl 2019, 13, (1), e1700179.

18. Guo, T.; Li, L.; Zhong, Q.; Rupp, N. J.; Charmpi, K.; Wong, C. E.; Wagner, U.; Rueschoff, J. H.; Jochum, W.; Fankhauser, C. D.; Saba, K.; Poyet, C.; Wild, P. J.; Aebersold, R.; Beyer, A., Multi-region proteome analysis quantifies spatial heterogeneity of prostate tissue biomarkers. Life Sci Alliance 2018, 1, (2).

19. Bache N1 G. P., 3, Bekker-Jensen DB3, Hoerning O1, Falkenby L1, Treit PV2, Doll S2, Paron I2, Müller JB2, Meier F2, Olsen JV3, Vorm O1, Mann M, A Novel LC System Embeds Analytes in Pre-formed Gradients for Rapid, Ultra-robust Proteomics. Mol Cell Proteomics. 2018.

20. Shi, J.; Wang, X.; Lyu, L.; Jiang, H.; Zhu, H. J., Comparison of protein expression between human livers and the hepatic cell lines HepG2, Hep3B, and Huh7 using SWATH and MRM-HR proteomics: Focusing on drug-metabolizing enzymes. Drug Metab Pharmacokinet 2018, 33, (2), 133–140.

21. He, B.; Shi, J.; Wang, X.; Jiang, H.; Zhu, H. J., Label-free absolute protein quantification with data-independent acquisition. J Proteomics 2019, 200, 51–59.

22. Colgrave, M. L.; Byrne, K.; Blundell, M.; Heidelberger, S.; Lane, C. S.; Tanner, G. J.; Howitt, C. A., Comparing Multiple Reaction Monitoring and Sequential Window Acquisition of All Theoretical Mass Spectra for the Relative Quantification of Barley Gluten in Selectively Bred Barley Lines. Anal Chem 2016, 88, (18), 9127–35.

23. Le Duff, M.; Gouju, J.; Jonchere, B.; Guillon, J.; Toutain, B.; Boissard, A.; Henry, C.; Guette, C.; Lelievre, E.; Coqueret, O., Regulation of senescence escape by the cdk4-EZH2-AP2M1 pathway in response to chemotherapy. Cell Death Dis 2018, 9, (2), 199.

24. Vowinckel, J.; Zelezniak, A.; Bruderer, R.; Mulleder, M.; Reiter, L.; Ralser, M., Cost-effective generation of precise label-free quantitative proteomes in high-throughput by microLC and data-independent acquisition. Sci Rep 2018, 8, (1), 4346.

25. Liu, Y.; Mi, Y.; Mueller, T.; Kreibich, S.; Williams, E. G.; Van Drogen, A.; Borel, C.; Frank, M.; Germain, P. L.; Bludau, I.; Mehnert, M.; Seifert, M.; Emmenlauer, M.; Sorg, I.; Bezrukov, F.; Bena, F. S.; Zhou, H.; Dehio, C.; Testa, G.; Saez-Rodriguez, J.; Antonarakis, S. E.; Hardt, W. D.; Aebersold, R., Multi-omic measurements of heterogeneity in HeLa cells across laboratories. Nat Biotechnol 2019, 37, (3), 314–322.

26. Yi Zhu 1, 3*, Tobias Weiss 4*, Qiushi Zhang 1,2, Rui Sun 1,2, Bo Wang 5, Zhicheng Wu 1,2, Qing Zhong 6,7, Xiao Yi 1,2, Huanhuan Gao 1,2, Xue Cai 1,2, Guan Ruan 1,2, Tiansheng Zhu 1,2, Chao Xu, Sai Lou 9, Xiaoyan Yu 10, Ludovic Gillet 3, Peter Blattmann 3, Karim Saba 11, Christian D. Fankhauser 11, Michael B. Schmid 11, Dorothea Rutishauser 6, Jelena Ljubicic 6, Ailsa, Christiansen 6, Christine Fritz 6, Niels J. Rupp 6, Cedric Poyet 11, Elisabeth Rushing 12, Michael Weller 4, Patrick Roth 4, Eugenia Haralambieva 6, Silvia Hofer 13, Chen Chen 14, Wolfram Jochum 15, Xiaofei Gao 1,2, Xiaodong Teng 5, Lirong Chen 10, Peter J. Wild 6,16#, Ruedi Aebersold 3,17#, Tiannan Guo, High-throughput proteomic analysis of FFPE tissue samples facilitates tumor stratification. biorxiv 2019.

27. Zhu, Y.; Guo, T., High-Throughput Proteomic Analysis of Fresh-Frozen Biopsy Tissue Samples Using Pressure Cycling Technology Coupled with SWATH Mass Spectrometry. Methods Mol Biol 2018, 1788, 279–287.

28. MacLean, B.; Tomazela, D. M.; Shulman, N.; Chambers, M.; Finney, G. L.; Frewen, B.; Kern, R.; Tabb, D. L.; Liebler, D. C.; MacCoss, M. J., Skyline: an open source document editor for creating and analyzing targeted proteomics experiments. Bioinformatics 2010, 26, (7), 966–8.

29. Rosenberger, G.; Koh, C. C.; Guo, T.; Rost, H. L.; Kouvonen, P.; Collins, B. C.; Heusel, M.; Liu, Y.; Caron, E.; Vichalkovski, A.; Faini, M.; Schubert, O. T.; Faridi, P.; Ebhardt, H. A.; Matondo, M.; Lam, H.; Bader, S. L.; Campbell, D. S.; Deutsch, E. W.; Moritz, R. L.; Tate, S.; Aebersold, R., A repository of assays to quantify 10,000 human proteins by SWATH-MS. Sci Data 2014, 1, 140031.

30. Rost, H. L.; Rosenberger, G.; Navarro, P.; Gillet, L.; Miladinovic, S. M.; Schubert, O. T.; Wolski, W.; Collins, B. C.; Malmstrom, J.; Malmstrom, L.; Aebersold, R., OpenSWATH enables automated, targeted analysis of data-independent acquisition MS data. Nat Biotechnol 2014, 32, (3), 219–23.

31. Goh, W. W. B.; Wong, L., Advanced bioinformatics methods for practical applications in proteomics. Brief Bioinform 2019, 20, (1), 347–355.

